# Energetic requirements and mechanistic plasticity in Msp1-mediated substrate extraction from lipid bilayers

**DOI:** 10.1101/2024.09.23.614443

**Authors:** Baylee Smith, Deepika Gaur, Nathan Walker, Isabella Walter, Matthew L. Wohlever

## Abstract

AAA+ proteins are essential molecular motors involved in numerous cellular processes, yet their mechanism of action in extracting membrane proteins from lipid bilayers remains poorly understood. One roadblock for mechanistic studies is the inability to generate subunit specific mutations within these hexameric proteins. Using the mitochondrial AAA+ protein Msp1 as a model, we created covalently linked dimers with varying combinations of wild type and catalytically inactive E193Q mutations. The wide range of ATPase rates in these constructs allows us to probe how Msp1 uses the energy from ATP hydrolysis to perform the thermodynamically unfavorable task of removing a transmembrane helix (TMH) from a lipid bilayer. Our *in vitro* and *in vivo* assays reveal a non-linear relationship between ATP hydrolysis and membrane protein extraction, suggesting a minimum ATP hydrolysis rate is required for effective TMH extraction. While structural data often supports a sequential clockwise/2-residue step (SC/2R) mechanism for ATP hydrolysis, our biochemical evidence suggests mechanistic plasticity in how Msp1 coordinates ATP hydrolysis between subunits, potentially allowing for robustness in processing challenging substrates. This study enhances our understanding of how Msp1 coordinates ATP hydrolysis to drive mechanical work and provides foundational insights about the minimum energetic requirements for TMH extraction and the coordination of ATP hydrolysis in AAA+ proteins.

## Introduction

Many fundamental cellular processes such as DNA replication, protein degradation, and vesicle trafficking require the physical remodeling of macromolecules. This energy intensive process is carried out by the ATPases Associated with diverse cellular Activities (AAA+) family of molecular motors which use the free energy of ATP binding and hydrolysis to perform mechanical work^1–6^. AAA+ proteins generally form hexameric rings and undergo ATP dependent movements to translocate a substrate through a narrow axial pore, resulting in substrate unfolding^7–12^.

Msp1 is homohexameric AAA+ ATPase anchored on the outer mitochondrial membrane (OMM) and peroxisome that promotes protein quality control by processively extracting mislocalized proteins or substrates that have become stalled in the Translocase of the Outer Membrane (TOM) complex^13–20^. The extracted substrates are then transferred to the ER where they are mostly ubiquitinated and degraded^21,22^. Loss of Msp1 results in severe mitochondrial stress including accumulation of mislocalized proteins, mitochondrial fragmentation, and loss of oxidative phosphorylation^23,24^. The human homolog ATAD1 has also been shown to disassemble the AMPA Receptor and regulate apoptosis by extracting the pro-apoptotic protein BIM^25,26^.

Understanding how ATP hydrolysis is coordinated between subunits is essential for understanding Msp1 function. Previous studies on other AAA+ unfoldases revealed that mechanical substrate unfolding is governed by a kinetic competition between substrate translocation and refolding of partially unfolded intermediates with robust substrate unfolding requiring several rapid rounds of ATP hydrolysis^27–31^. It is unclear if a similar kinetic competition between TMH extraction and re-insertion also governs Msp1 activity.

A major obstacle for studying the coordination of ATP hydrolysis in Msp1 homohexamers is the inability to generate subunit specific mutations. Drawing inspiration from previous work on the AAA+ ATPase ClpX^32^, we sought to overcome this challenge by using genetically encoded flexible linkers to create covalent Msp1 dimers **(Figure 1A)**. The resulting constructs form a trimer of dimers rather than a standard hexamer composed of six monomers, thereby allowing the generation of subunit specific mutations.

**Figure 1:**
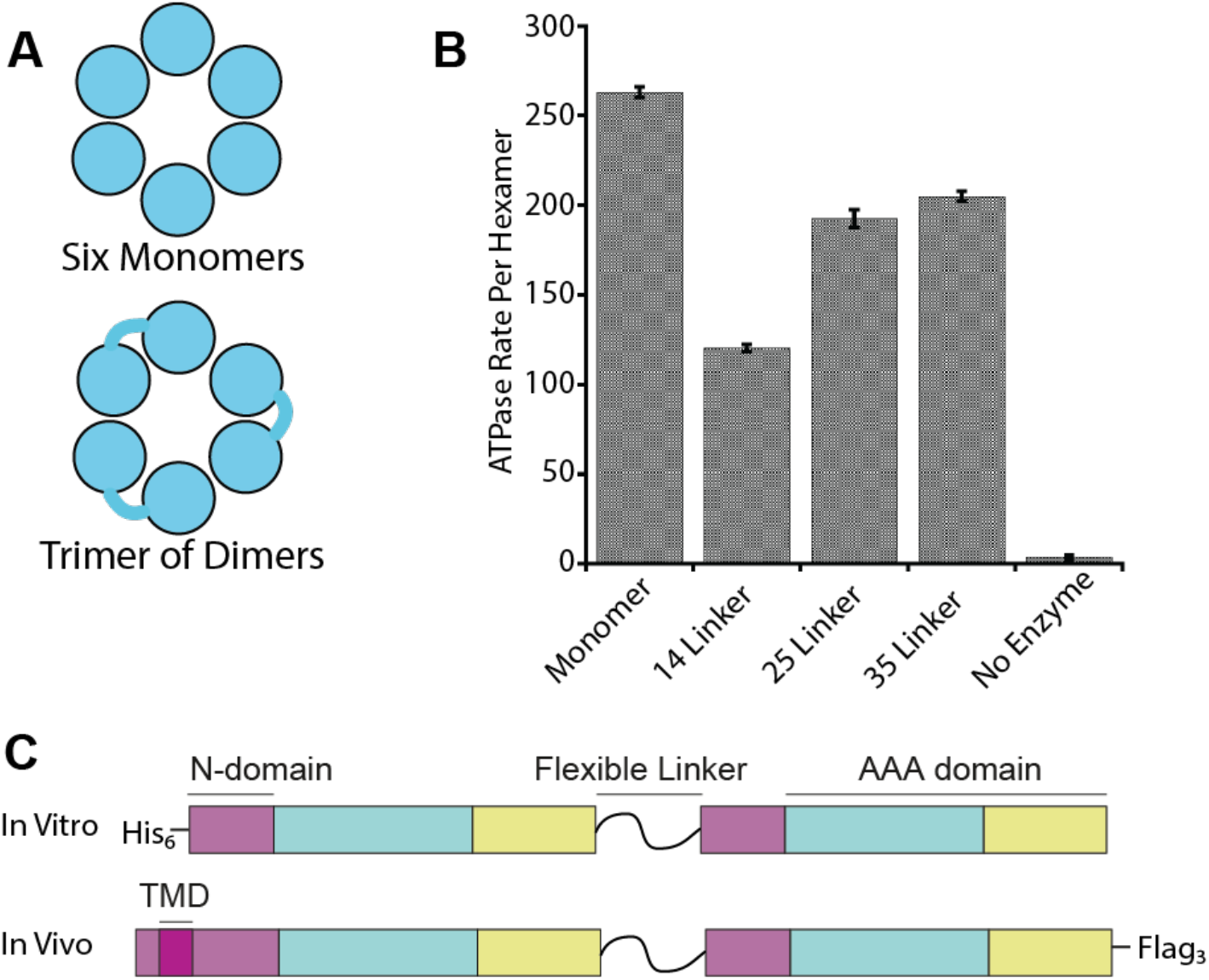
Design of Covalently Linked Dimers. A) Diagram showing how covalently linked dimers allow for the design of subunit specific mutations by forming a trimer of dimers. B) ATPase assay shows that dimers with a linker length of 35 residues are most active. C) Diagram showing the design of the covalently linked dimers for use *in vitro* and *in vivo*.

Here, we developed covalently linked Msp1 dimers containing varying combinations of WT and ATPase deficient Walker B mutant (E193Q) subunits. These constructs have different maximum rates of ATP hydrolysis and thus serve as an allelic series that allows us to probe the ATP hydrolysis requirements for Msp1 functionality both *in vitro* and *in vivo*. Our results demonstrate that hexamers with up to 50% inactive subunits retain robust activity, suggesting that Msp1 has mechanistic plasticity to bypass inactive subunits. Interestingly, the overall level of functionality depends on the type of activity being measured, suggesting that different aspects of Msp1 function require different minimum rates of ATP hydrolysis. Overall, our results provide foundational insights into how Msp1 coordinates ATP hydrolysis between six subunits to overcome the substantial thermodynamic barriers that accompany substrate extraction from a lipid bilayer.

## Results

### Development of covalently linked dimers

A major obstacle in the development of linked dimers is the design of a linker that will not significantly impact the function of the enzyme. The genetically encoded linker needs to have sufficient length and flexibility to minimize disruption of catalytic activity. Based on the cryo-EM structure of soluble Msp1 (PDB = 6PDW), the distance between N and C-termini in adjacent subunits ranges from 55 to 76 Å^33^. In principle the shortest distance could be bridged by 16 fully-extended amino acids whereas the longest distance would require more than 22 amino acids. These are only rough estimates as the termini are not visible in the structure and any linker between subunits would need to adopt a longer, more flexible conformation to avoid steric clashes.

To determine the optimal linker length, we generated three different soluble constructs consisting of two wild-type subunits with linkers of 14, 25, or 35 amino acids. We measured ATPase rates as a proxy for how the linker lengths affected Msp1 activity. We observed that constructs with a 35 amino acid linker had the smallest decrease in ATPase activity compared to the WT monomer **(Figure 1B)**. As the 35 amino acid linker can accommodate even the longest possible distance between subunits, we proceeded with a 35 amino acid linker.

The final version of the linked Msp1 constructs were generated by gene synthesis to allow for each subunit have identical protein sequences but unique DNA sequences. The linker is composed of small, hydrophilic amino acids and contains a unique restriction enzyme site, allowing for individual subunits to be replaced with mutated versions via restriction cloning.

We generated two versions of the linked dimers, a soluble version for *in vitro* assays and a full-length version for *in vivo* assays **(Figure 1C)**. Soluble constructs have a deletion of the first 32 amino acids in each subunit. There is an N-terminal His_6_ tag to aid in substrate purification and the *in vitro* extraction assay. For *in vivo* assays we generated “full-length” constructs where the first subunit contains the native TMD and second subunit has a deletion of the first 32 amino acids to remove the TMD. To allow detection by western blot, we added a 3x Flag tag at the C-terminus.

### Linked constructs retain activity *in vivo*

To better understand how Msp1 couples ATP hydrolysis to mechanical work, we generated three constructs containing a mixture of wild type (WT) and E193Q (EQ) Walker B mutations. The Walker B mutation allows ATP to bind to a subunit but prevents hydrolysis. We refer to these constructs as WT-WT, WT-EQ, and EQ-WT.

To test if the linked dimers are active *in vivo*, we cloned the full-length version into a centromeric plasmid with the native Msp1 promoter. Anti-Flag western blots show relatively equal expression across the three dimers **(Figure 2A)**. The dimer bands have lower signal intensity than monomeric Msp1. However, only three copies of the dimers are required to form a functional hexamer rather than six copies of the monomer. Thus, the overall concentration of hexamers is roughly comparable between the monomeric and dimeric constructs.

**Figure 2:**
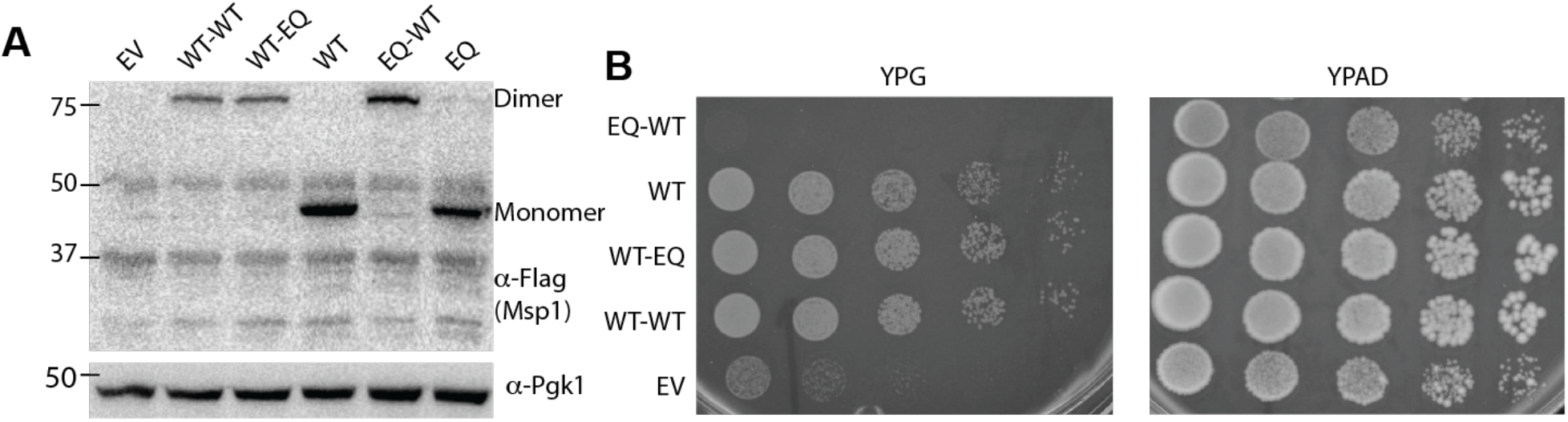
Several linked constructs retain activity *in vivo*. A) Anti-Flag western blot shows roughly equal expression of linked dimers. B) Complementation assay shows that WT-WT and WT-EQ can rescue growth of *Msp1Δ, Get3Δ* cells on glycerol, whereas EQ-WT cannot.

As a first test of Msp1 activity *in vivo* we performed a complementation assay. Simultaneous deletion of *Msp1* and *Get3* leads to a loss of oxidative phosphorylation which can be assayed by growth on the non-fermentable carbon source glycerol^17^. We therefore generated *Msp1Δ, Get3Δ* yeast strains complemented with a centromeric plasmid. As expected, complementation with the empty vector failed to rescue growth on glycerol whereas plasmid-based expression of WT Msp1 provided a complete rescue. Surprisingly, complementation with both WT-WT and WT-EQ provided a complete rescue, implying that Msp1 does not require maximum ATP hydrolysis rates to be functional *in vivo*. The EQ-WT construct provided no rescue and appeared to have a slight toxic effect compared to the empty vector control **(Figure 2B)**. We conclude that the WT-WT and WT-EQ constructs retain functionality *in vivo* while the EQ-WT construct is inactive.

### The WT-EQ construct has fully active WT subunits

We next asked why there was a difference in activity between the WT-EQ and EQ-WT constructs despite both constructs containing a 50-50 mixture of WT and EQ subunits. As Msp1 activity depends on ATP hydrolysis, we hypothesized that the WT-EQ construct had a higher rate of ATP hydrolysis than the EQ-WT construct. ATPase measurements confirmed our hypothesis **(Figure 3A)**. While the EQ-WT construct had a lower rate of ATP hydrolysis, it still retained activity, suggesting that a minimum rate of ATP hydrolysis is required for Msp1 function *in vivo*.

**Figure 3:**
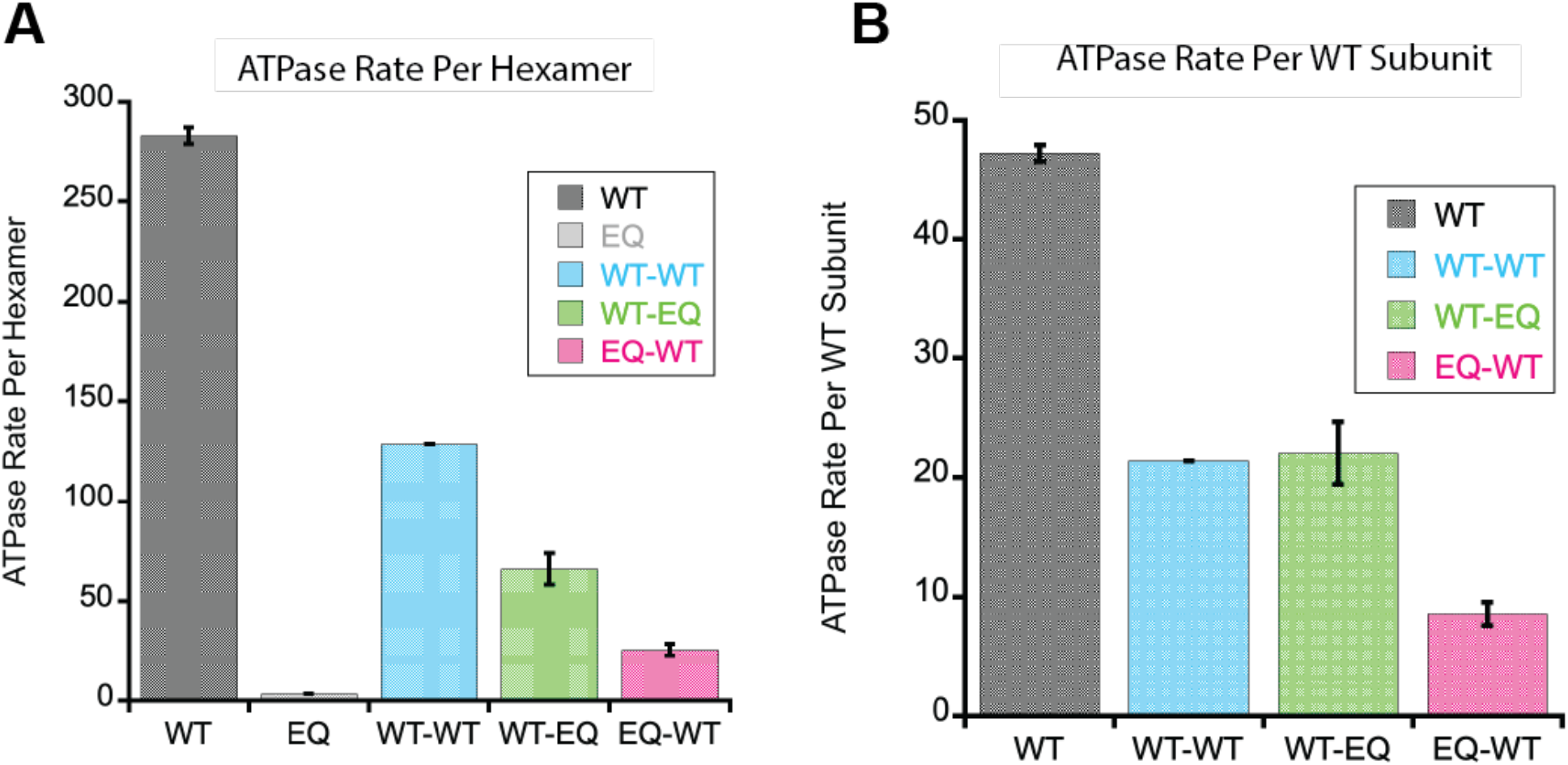
The WT-EQ construct has fully active WT subunits. A) ATPase activity of linked dimers normalized per hexamer. The WT-EQ construct has higher ATPase activity than the EQ-WT construct. Note that there are slight differences in construct design and purification buffers compared to the initial screening in Figure 1, which can account for the differences in activity. B) As in A, but ATPase activity of linked dimers is normalized per WT subunit. There is no difference in ATPase activity of WT subunits between WT-WT and WT-EQ constructs.

Interestingly, when the ATPase rate is normalized to the number of WT subunits, we observed no difference in the rate of ATP hydrolysis between WT-WT and WT-EQ **(Figure 3B)**. This indicates that the neighboring EQ subunit in this construct has no adverse effect on ATPase activity in the WT subunit.

### *In vivo* extraction of Pex15ΔC30 correlates with ATPase rates

While ATP hydrolysis is clearly essential for Msp1 function, our current understanding of how the rate of ATP hydrolysis affects different aspects of Msp1 function are limited. We reasoned that our linked dimers could function as an allelic series to determine the ATP hydrolysis requirements for Msp1 activity.

We first asked what is required for substrate extraction *in vivo*. To measure Msp1 substrate extraction activity *in vivo* we used a GFP/mCherry reporter assay^34^. Briefly, GFP and mCherry are expressed on a polycistronic vector separated by the codon skipping E2A site, giving rise to equal expression of GFP and mCherry in the cell **(Figure 4A)**. We then fused the known Msp1 substrate Pex15ΔC30 to the C-terminus of GFP. The ΔC30 truncation causes Pex15 to constitutively mislocalize to the mitochondria^18^, where it is extracted by Msp1 and eventually degraded^21^. Msp1 activity is measured by the ratio of GFP to mCherry in cells via flow cytometry, with a lower GFP/mCherry ratio indicating higher levels of Msp1 activity. Control reactions with *Msp1Δ, Get3Δ* yeast complemented with WT Msp1 on a centromeric plasmid show that the GFP/mCherry ratio is ∼1 with untagged GFP and decreases to ∼0.5 when Pex15ΔC30 is fused to GFP **(Figure 4B)**.

**Figure 4:**
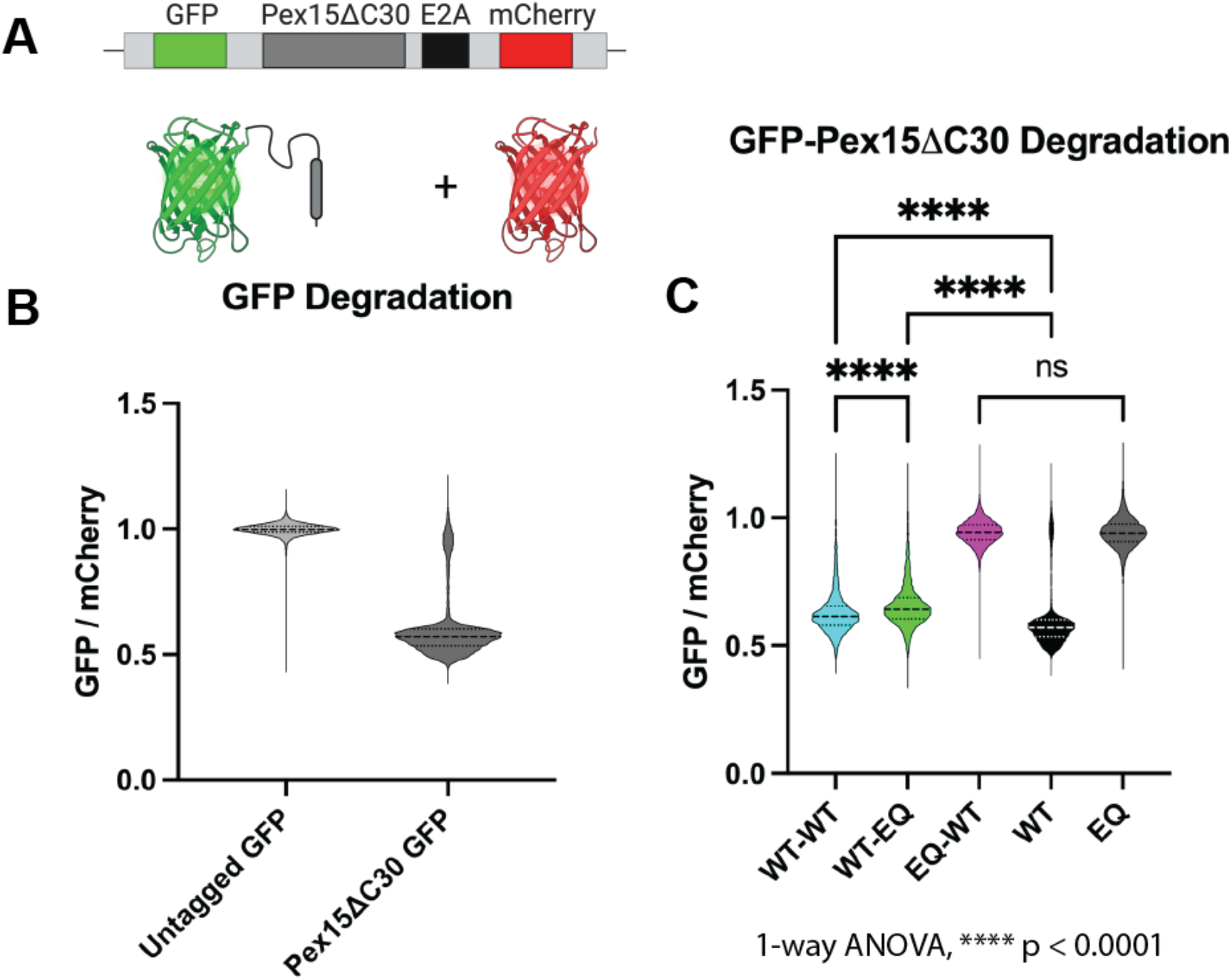
In vivo extraction of Pex15ΔC30 correlates with ATPase rates. A) Diagram of the reporter used for the *in vivo* extraction assay. B) Control reactions show that degradation of GFP depends on Pex15ΔC30. *C) In vivo* extraction assays shows modest but significant differences in extraction activity of WT, WT-WT, and WT-EQ constructs that correlate with ATPase rates. Despite retaining ATPase activity, the extraction activity of the EQ-WT construct is indistinguishable from the EQ construct.

We then used this assay to test the activity of the linked dimers **(Figure 4C)**. Consistent with the complementation assay, the EQ-WT construct was indistinguishable from the negative control EQ monomer. The WT monomer, WT-WT, and WT-EQ constructs showed robust activity. Importantly, the trend parallels the ATPase rates with WT monomer being the most active, WT-EQ being the least active, and WT-WT falling between the two.

### There is a non-linear correlation between substrate extraction and ATPase rate

Classical studies on other AAA+ unfoldases demonstrate that there is a non-linear relationship between ATPase activity and substrate unfolding^27^. AAA+ proteins that process membrane proteins must overcome the additional thermodynamic barrier of removing a TMH from a lipid bilayer^35,36^. To test if there is a similar relationship between ATPase rates and substrate extraction, we turned to our *in vitro* extraction assay^37^.

The *in vitro* extraction assay utilizes a split-luciferase system, with substrate extraction monitored by luminescence^35^. We performed the extraction assay with the Sec22 model substrate and standard mitochondrial liposomes, which contain a mixture of lipids meant to mimic the mitochondrial membrane. The WT-WT construct had activity levels close to the WT monomer. Surprisingly all other constructs, including the WT-EQ construct were essentially inactive **(Figure 5A)**, suggesting that the *in vitro* assay is a more stringent test of Msp1 activity.

**Figure 5:**
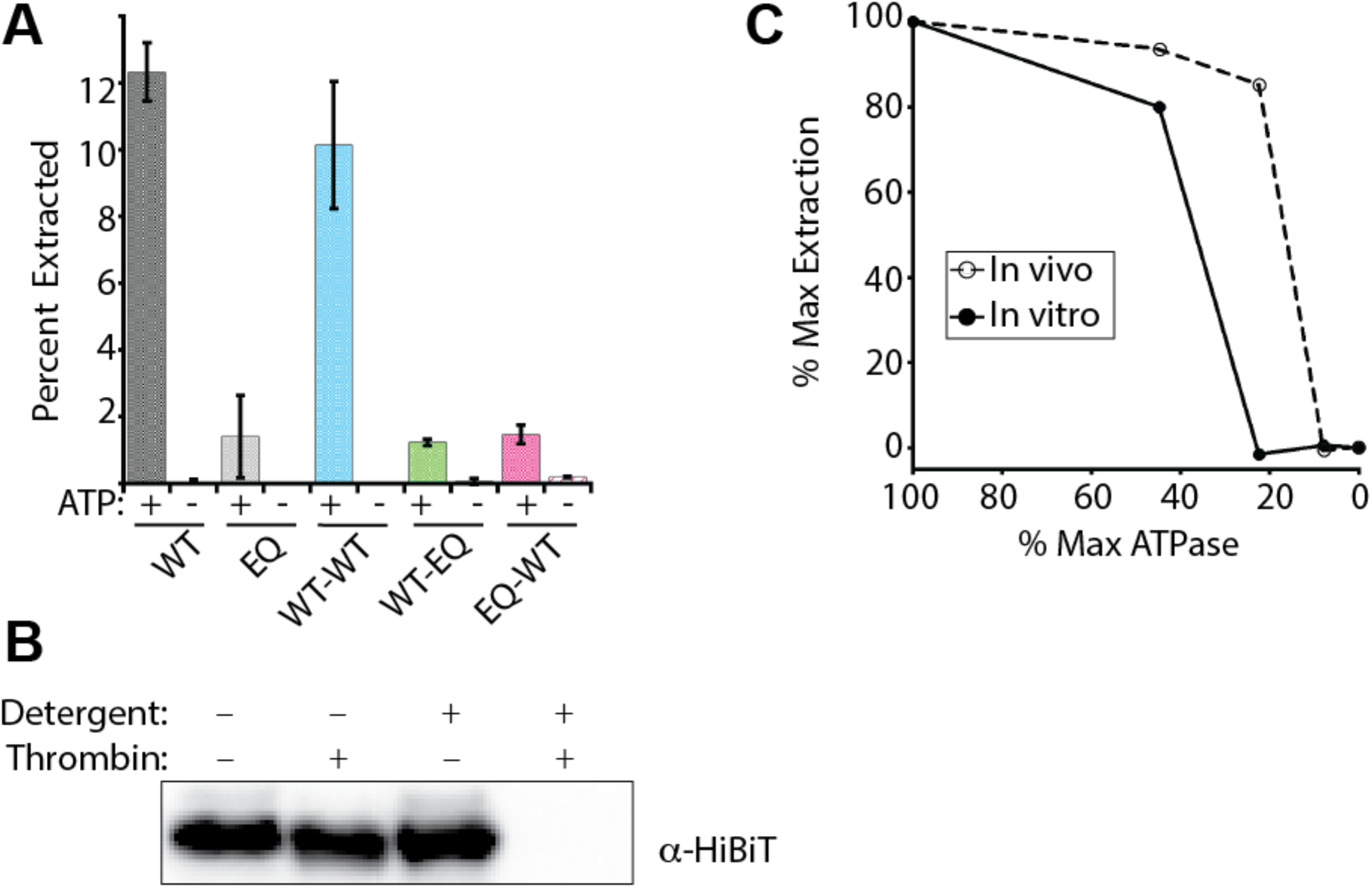
There is a non-linear correlation between substrate extraction and ATPase rate. A) Extraction assay of the Sec22 model substrate in standard liposomes shows that WT-WT retains > 80% extraction activity whereas the EQ-WT and WT-EQ constructs are inactive. B) Protease protection assay demonstrates that the Sec22 model substrate is properly oriented in the standard liposomes. A thrombin protease site is engineered into the substrate between the TMD and HiBiT tag. Properly oriented substrate has the thrombin site sequestered in the lumen of the liposomes, where it is only accessible upon detergent solubilization. Thrombin cleavage results in a 14 residue HiBiT fragment, which is not detectable by western blot. C) Rates of substrate extraction vs ATPase rate show a non-linear relationship. *In vivo* extraction of Pex15 (open circles) is the average GFP/mCherry ratio from Figure 4C and *in vitro* extraction data (closed circles) is from Figure 5B. Activity is normalized such that WT and EQ correspond to 100% and 0% activity, respectively.

## Discussion

AAA+ proteins are ubiquitous molecular motors that are involved in numerous essential cellular processes. These hexameric proteins convert ATP hydrolysis to mechanical work, resulting in substrate remodeling. While significant effort has been devoted to understanding how AAA+ proteins drive protein unfolding, comparatively little is known about how AAA+ proteins extract membrane proteins from the lipid bilayer. Here, we used the mitochondrial AAA+ protein Msp1 as a model system to examine this important topic. We generated covalently linked Msp1 dimers containing a mixture of wild type and ATPase deficient Walker B mutants that are active both *in vitro* and *in vivo*. The resulting trimers of dimers have varying rates of ATP hydrolysis and thus serve as an allelic series that we used to probe how ATP hydrolysis is coordinated between Msp1 subunits for TMH extraction from the lipid bilayer.

Using a combination of *in vitro* and *in vivo* assays, we demonstrate that there is a non-linear relationship between ATP hydrolysis and Msp1 extraction activity **(Figure 5C)**. For example, the WT-WT construct has only 45% the ATPase activity of the WT monomer but retains 80% and 93% extraction activity *in vivo* and *in vitro*. Conversely, the EQ-WT construct has 8% of the ATPase activity of the WT monomer and is inactive in all assays. Interestingly, the WT-EQ construct, which has 22% ATPase activity, is robustly active *in vivo* and inactive for the *in vitro* extraction assay. The non-linear relationship between ATP hydrolysis and substrate extraction suggests that a minimum rate of ATP hydrolysis is required to extract a TMH from a lipid bilayer, with the minimum rate of extraction depending on the substrate and assay.

Interestingly, all assays suggest that the rate of ATP hydrolysis by Msp1 far exceeds the minimum ATPase rate required for activity. Much like a typical car does not utilize all available horsepower when cruising on the highway, we propose that the “reserve horsepower” in Msp1 may be important for extracting particularly challenging substrates which are not captured in our assays.

Consistent with our results, a non-linear relationship between ATPase rates and substrate processing was previously observed with the AAA+ protein ClpX^27^. This was ascribed to the stepwise manner of substrate processing leading to a partially unfolded intermediate.

Competition between unfolding and refolding of the intermediate is governed by the rate of ATP hydrolysis. In this model, rapid ATP hydrolysis favors complete substrate unfolding and slow ATP hydrolysis favors substrate refolding. We propose that a similar relationship exists with Msp1-mediated TMH extraction from a lipid bilayer. High resolution assays, ideally with real-time kinetics, will be required to definitively test this hypothesis.

The relationship between ATP hydrolysis and substrate processing intimately depends on the coordination of ATP hydrolysis between subunits in the Msp1 homohexamer. Recent cryo-EM structures of *Chaetomium thermophilum* Msp1 and human ATAD1 show a canonical right-handed spiral arrangement of subunits that has been observed in many other AAA+ proteins^33,38^. While the exact details vary depending on the particular combination of ATP analog and ATPase deficient mutants used, these structures generally resemble a lock washer with five ATP/ADP bound subunits arranged in a spiral staircase and a transitional sixth subunit in the apo state. The substrate is gripped in the central pore by highly conserved loops, referred to as pore loops, which intercalate between the side chains of the substrate. The pore loops are arranged like a spiral staircase with a two amino acid step size. These structures suggest that ATP hydrolysis occurs in an ordered sequence around the ring. As subunits proceed through the steps of ATP binding, hydrolysis, and release, they transition up the spiral, thereby translocating the substrate. This mechanism has been termed the sequential clockwise/2-residue step (SC/2R) mechanism^8,39^.

One prediction of this model is that the presence of a catalytically inactive subunit should disrupt the highly ordered sequence of ATP hydrolysis, acting in a dominant negative fashion to poison the ring. We provide multiple lines of evidence that Msp1 hexamers containing multiple EQ subunits retain activity *in vitro* and *in vivo* **(Figures 2-5)**, which is inconsistent with the SC/2R mechanism.

While cryo-EM structures of diverse AAA+ proteins almost universally support the SC/2R mechanism, biochemical experiments suggest that there is likely more mechanistic plasticity than are revealed by these static structures^12,40,41^. For example, single turnover experiments with the AAA+ protein ClpA showed a step size of 14 amino acids alone or 5 amino acids when in complex with the ClpP Protease^42,43^. Likewise, covalently linked ClpX hexamers containing multiple ATPase inactive subunits are still capable of unfolding and translocating substrates^32^. Single molecule optical trapping experiments with ClpXP showed a basic step size of 5-8 to 10-13 amino acids^31^. It is possible that the larger step sizes are actually composed of multiple unresolved SC/2R steps, but the differing kinetics of substrate translocation and ATP hydrolysis argue against this possibility^39^. The inconsistency between biochemical and structural data has given rise to other mechanisms for coordinating ATP hydrolysis between subunits, such as the Probabilistic Anti-clockwise Long-Step (PA-LS) or Sequential Clockwise/6-residue (SC/6R)^12,39^.

While further work is needed to determine the details for how Msp1 subunits coordinate ATP hydrolysis, we conclude that Msp1 does not operate solely by the SC/2R mechanism of ATP hydrolysis, but instead shows mechanistic plasticity. We propose that this mechanistic plasticity may provide robustness to prevent stalling during the translocation of challenging substrates, similar to mechanisms used by soluble AAA+ proteins^40,44–47^.

Overall, our work sheds light on how Msp1 utilizes ATP hydrolysis to perform mechanical work both *in vitro* and *in vivo*. It also raises interesting questions regarding the minimum rates of ATP hydrolysis required to overcome the thermodynamic barrier of TMH extraction from a lipid bilayer and how Msp1 coordinates ATP hydrolysis between subunits. The Msp1 dimers presented here will be a powerful tool for addressing these important questions.

## Materials and Methods

### Msp1 construct design and cloning

Soluble Msp1 dimers were restriction cloned with NdeI and XhoI into a pET28a vector containing an N-terminal His_6_ tag followed by a TEV protease site. Dimers were generated by gene synthesis such that each subunit has the same protein sequence but a unique DNA sequence. The linker sequence is ASGSGGSEASASAGAAGSGDGSGSGGSEGGTSGAT and contains a HindIII restriction cloning site. To generate point mutations, the individual subunits were first subcloned into a pET28a vector, mutated by Quikchange PCR, and then restriction cloned back into the dimer vector.

### Protein expression and purification

#### Protein expression

Briefly, constructs were transformed in BL21 pRIL cells, grown in terrific broth at 37° C until OD_600_ = ∼0.6-0.8. Cells were induced with 250 µM isopropyl β-D-1-thiogalactopyranoside (IPTG) for 4 hours at either 20° C for Msp1 constructs, overnight at 16° C for chaperone constructs, overnight at 18° C for the Sec22 substrate, or for 4 hours at 25° C for all other constructs. Cells were pelleted by centrifugation, resuspended in appropriate Lysis Buffer supplemented with 0.05 mg/mL Lysozyme, 1 mM Protease Inhibitor (GoldBio), and 1 mM phenylmethylsulfonyl fluoride (PMSF) and stored at -80° C. Msp1 pellets were also supplemented with 1 mM Protease Inhibitor (GoldBio) prior to storage at -80° C. For purification, the cell pellet was rapidly thawed, supplemented with 500 U of universal nuclease (Pierce), lysed by sonication, and the supernatant was isolated by centrifugation for 30 minutes at 18,000 x g.

#### Msp1 constructs

Prior to sonication, Msp1 constructs were supplemented with additional Lysis Buffer containing a final concentration of 1 mM ATP and 1 mM PMSF. Supernatant for Msp1 constructs were loaded onto a gravity column with Ni-NTA resin (Qiagen) and washed with 20 column volumes (CV) of Msp1 Lysis Buffer (20 mM Tris pH 7.5, 200 mM potassium acetate, 20 mM imidazole, 1 mM DTT, 0.01 mM EDTA) followed by 10 CV of Msp1 Wash Buffer (20 mM Tris pH 7.5, 100 mM potassium acetate, 30 mM imidazole, 1 mM DTT, 0.01 mM EDTA). Samples were eluted with 2.5 CV of Msp1 Elution Buffer (20 mM Tris pH 7.5, 100 mM potassium acetate, 250 mM imidazole, 1 mM DTT, 0.01 mM EDTA, 1 mM ATP). Samples were spin concentrated to 3-10 mg/mL in a Amicon Ultra centrifugal filter (Pierce) and further purified by size exclusion chromatography with a Superdex200 Increase 10/300 GL column equilibrated in Msp1 FPLC Buffer (20 mM Tris pH 7.5, 200 mM potassium acetate,1 mM DTT). Pure fractions as judged by SDS PAGE were pooled, spin concentrated to 3-5 mg/mL in an Amicon Ultra centrifugal filter (Pierce), aliquoted, and flash frozen in single use aliquots. Protein concentration was measured by Bradford Assay.

#### SGTA and calmodulin

Lysates for GST-SGTA and GST-calmodulin were loaded onto a gravity column containing glutathionine resin (Thermo Fisher). Resin was washed with 15 CV of SGTA Lysis Buffer (50 mM Hepes-KOH pH 7.5, 150 mM NaCl, 1 mM DTT, 0.01 mM EDTA, 10% glycerol). Protein was eluted with 3 CV of SGTA Lysis Buffer supplemented with 10 mM reduced glutathione. The protein was further purified by size exclusion chromatography with a Superdex200 Increase 10/300 GL column equilibrated in SGTA FPLC Buffer (20 mM Hepes-KOH pH 7.5, 100 mM NaCl, 0.1 mM TCEP). Pure fractions as judged by SDS PAGE were pooled, spin concentrated to 15-30 mg/mL in an Amicon Ultra centrifugal filter (Pierce), aliquoted, and flash frozen in single use aliquots. Protein concentration was measured by absorbance at 280 nm using the extinction coefficient from Expasy Protparam^48^.

#### LgBiT and MBP-Ubiquitin-mNG2

LgBiT and MBP-Ubiquitin-mNG2 were purified by Ni-NTA affinity chromatography. Cleared lysate was loaded onto a gravity column containing Ni-NTA resin (Qiagen) and washed with 15 CV of LgBiT Lysis Buffer (20 mM Tris pH 7.5, 200 mM NaCl, 20 mM imidazole, 1 mM DTT, 0.01 mM EDTA) followed by 5 CV of LgBiT Wash buffer (20 mM Tris pH 7.5, 200 mM NaCl, 50 mM imidazole, 1 mM DTT, 0.01 mM EDTA). Samples were eluted with LgBiT Elution Buffer (20 mM Tris pH 7.5, 200 mM NaCl, 250 mM imidazole, 1 mM DTT, 0.01 mM EDTA) and spin concentrated to 5-10 mg/mL in an Amicon Ultra centrifugal filter (Pierce). LgBiT was cleaved with 0.01 mg/mg ratio of 3C protease to substrate overnight at 4° C. Samples were further purified by size exclusion chromatography with a Superdex200 Increase 10/300 GL column equilibrated in LgBiT FPLC Buffer (20 mM Tris pH 7.5, 200 mM NaCl, 1 mM DTT). Pure fractions as judged by SDS PAGE were pooled, spin concentrated to 10-15 mg/mL in an Amicon Ultra centrifugal filter (Pierce), aliquoted, and flash frozen in single use aliquots. Protein concentration was measured by absorbance at 280 nm using the extinction coefficient from Expasy Protparam^48^.

#### SumoTMD substrate

The SumoTMD substrate was expressed, harvested and lysed as described above using the SumoTMD Lysis Buffer (50 mM Tris pH 7.5, 300 mM NaCl, 10 mM MgCl_2_, 10 mM imidazole, 10% glycerol. After sonication, membrane proteins were solubilized by the addition of n-dodecyl--Δ-D-maltoside (DDM) to a final concentration of 1% and rocked at 4° C for 30 minutes. Lysate was cleared by centrifugation at 35,000 x g for 1 h. Cleared lysate was loaded onto a gravity column with Ni-NTA resin (Qiagen) and washed with 5 CV of SumoTMD Wash Buffer 1 (50 mM Tris pH 7.4, 500 mM NaCl, 10 mM MgCl_2_, 10 mM imidazole, 5 mM Δ-mercaptoethanol (BME), 10% glycerol, 0.1% DDM). Resin was washed with 5 CV of SumoTMD Wash Buffer 2 (50 mM Tris pH 7.4, 300 mM NaCl, 10 mM MgCl_2_, 25 mM imidazole, 5 mM BME, 10% glycerol, 0.1% DDM) and 5 CV of SumoTMD Wash Buffer 3 (50 mM Tris pH 7.4, 150 mM NaCl, 10 mM MgCl_2_, 50 mM imidazole, 5 mM BME, 10% glycerol, 0.1% DDM). Sample was then eluted with 2 CV of SumoTMD Elution Buffer (50 mM Tris pH 7.5, 150 mM NaCl, 10 mM MgCl_2_, 250 mM imidazole, 5 mM BME, 10% glycerol, 0.1% DDM). Samples were spin concentrated in an Amicon Ultra centrifugal filter (Pierce) and cleaved with 0.01 mg/mg ratio of 3C protease/substrate overnight.

On day 2, the sample volume was brought up to 5 mL using SumoTMD FPLC Buffer (50 mM Tris pH 7.4, 150 mM NaCl, 10 mM MgCl_2_, 5 mM BME, 10% glycerol, 0.1% DDM). To remove uncleaved protein and 3C protease, samples were incubated with Ni-NTA resin for 30 minutes and loaded onto a column. The flow through was collected and spin concentrated. Samples were further purified by size exclusion chromatography with a Superdex200 Increase 10/300 GL column equilibrated in SumoTMD FPLC Buffer. Pure fractions as judged by SDS PAGE were pooled, spin concentrated to 5 mg/mL in an Amicon Ultra centrifugal filter (Pierce), aliquoted, and flash frozen in single use aliquots. Protein concentration was measured by absorbance at 280 nm using the extinction coefficient from Expasy Protparam^48^.

### ATPase assays

ATPase activity was determined using a coupled ATPase assay modified from previous work on ATPases^49^. Reactions were setup in a 96-well clear bottom plate. Each reaction contained a final concentration of 50 mM Hepes-KOH (pH 7.5), 200 mM potassium acetate, 2 mM DTT, 1 mM Phosphoenolpyruvate, 0.3 mM NADH, 20 U/mL Lacate Dehydrogenase, 10 U/mL Pyruvate Kinase, 2 mM ATP, and 1 µM of the desired Msp1 construct normalized to hexamer concentration. Reactions were performed in triplicate. Plate was incubated at 30° C for 15 minutes. Reaction was initiated by the addition of 100 mM magnesium acetate to a final concentration of 10 mM. Absorbance at 340 nm was read every 11 seconds for a 10-minute period.

### Liposome preparation and reconstitution

Liposomes were prepared as described previous^37^. Liposomes mimicking the OMM were prepared by mixing chloroform stocks of Chicken egg phosphatidyl choline (Avanti 840051C), chicken egg phosphatidyl ethanolamine (Avanti 840021C), bovine liver phosphatidyl inositol (Avanti 840042C), synthetic DOPS (Avanti 840035C), synthetic TOCL (Avanti 710335C), and 1,2-dioleoyl-sn-glycero-3-[N-((5-amino-1-carboxypentyl)iminodiacetic acid)succinyl] Nickel salt (Avanti 790404) at a 48:28:10:8:4:2 molar ratio with 1 mg of DTT.

Chloroform was evaporated under a gentle stream of nitrogen and then left on a vacuum (<1 mTorr) overnight. Lipid film was fully resuspended in Liposomes Buffer (50 mM Hepes KOH pH 7.5, 15 % glycerol, 1 mM DTT) over the course of several hours by vortexing and rotation on a wheel at room temperature. The final concentration of liposomes was 20 mg/mL. The liposomes were subjected to five freeze-thaw cycles with liquid nitrogen and rapid thawing followed by 15x extrusion through a 100 nm filter at 60° C. Single use aliquots were flash frozen in liquid nitrogen and stored at -80° C.

SumoTMD construct was reconstituted into liposomes by mixing 2.5 μM SumoTMD and 2 mg/mL Liposomes. The final volume was brought to 100 μL with Reconstitution Buffer (50 mM Hepes-KOH pH 7.5, 200 mM potassium acetate, 7 mM magnesium acetate, 2 mM DTT, 10% sucrose 0.01% sodium azide, and DeoxyBigChap (DBC)). The concentration of DBC used for reconstitution was re-optimized for each liposome preparation but ranged from 0.1%-0.5%.

Detergent was removed by adding 25 mg of biobeads and rotating the samples for 16 hours at 4° C. After removing the biobeads, the reconstituted material was incubated with 5 μM GST-SGTA and 5 μM GST-Calmodulin to remove unincorporated material. The sample was diluted with 100 μL Extraction Buffer (50 mM Hepes-KOH pH 7.5, 200 mM potassium acetate, 7 mM magnesium acetate, 2 mM DTT, 0.1 μM CaCl_2_). Samples were incubated with glutathione spin columns (Pierce) for 30 minutes. Flow through material was collected and used as pre-cleared material for the extraction assay.

Extraction reactions were setup using pre-cleared material with a final concentration of 3 μM Msp1 construct normalized per hexamer, 1 mg/mL bovine serum albumin (BSA), 5 µM SGTA, 5 µM calmodulin, and 0.085 µM CaCl_2_. Reactions were brought up to final volume with extraction buffer lacking CaCl_2_ and were performed in triplicate. Reaction tubes were incubated at 30° C for 1-2 minutes before addition of 80 mM ATP. Reactions were then incubated at 30° C for 25 minutes before the addition of 8 µM MBP-Ubiquitin-mNG2. They were incubated for another 5 minutes before transfer to TLA-120.1 centrifuge tubes. Samples were pelleted at 65,000 rpm for 30 minutes. 20 µL of supernatant was taken from the top of the samples following centrifugation. These samples were incubated with 1.5 µM LgBiT. Samples were brought up to a total volume of 50 µL with extraction buffer. Following incubation, samples and full signal controls were transferred to individual wells of a white half-volume 96-well plate. 20 µL of Promega furimazine was added to each well. Luminescence was read at 470 nm with a 1 mm read height with a 1 second integration every 32 seconds for 10 minutes. Peak luminescence values were used for calculation of percent signal.

### Protease protection assay

A protease protection assay was performed on pre-clear material used for extraction assays. The final reaction had a total volume of 10 µL and contained 7 µL of pre-cleared material, 2 U of thrombin protease, and 1% of Triton X-100 (where indicated). Samples were incubated at room temperature for 1 hour and then 1 µL of 200 mM PMSF was added. The samples were reverse quenched into 90 µL of boiling 1% sodium dodecyl sulfate and incubated at 95° C for 10 minutes. Anti-HiBiT western blot was performed with 1:5,000 dilution of mouse anti-HiBiT (Promega, Clone 30E5) and 1:10,000 dilution of goat anti-mouse-HRP secondary antibody (Invitrogen).

### Yeast complementation assay

Strains and plasmids used in the study are described in **Tables 1** and **2**. For complementation assays in *S. cerevisiae, Msp1Δ, Get3Δ* W303-1 yeast were generated by integrating Msp1::KanMX and Get3::NatMX using homologous recombination in WT 303-1 strain. Msp1 constructs were cloned into a *CEN* and *LEU2* vector with the flanking 281 bp upstream and 261 bp downstream as the respective promoter and terminator. Msp1 construct expression in *S*.

**Table 1:**
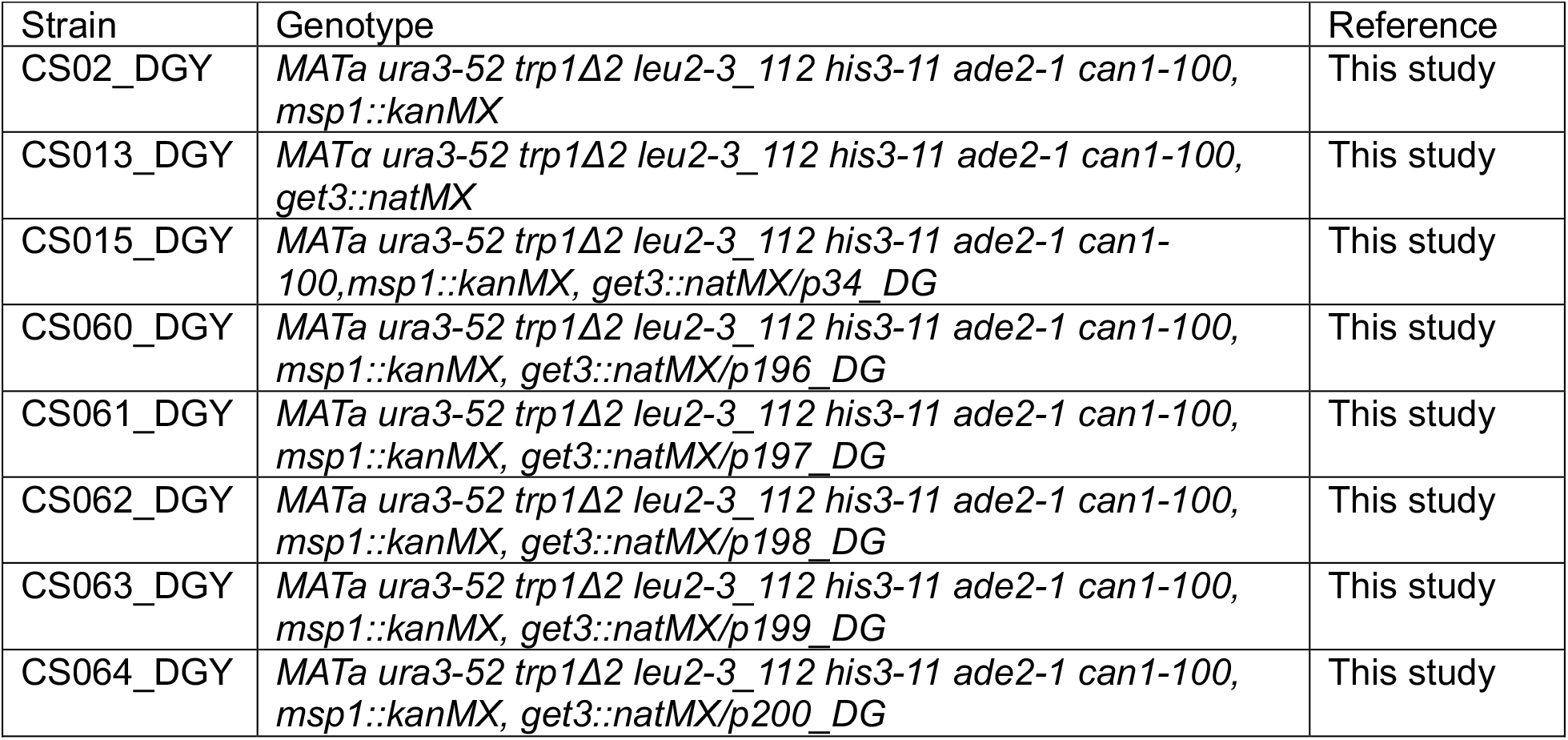
Strains used in this study.

**Table 2:** Plasmid used in this study.

| Plasmid | Description | Reference |
| --- | --- | --- |
| p34_DG | *pRS315* |  |
| p196_DG | *CEN, LEU2, P<sub>MSP1</sub>-MSP1(WT)-MSP1(WT)-3XFLAG-T<sub>MSP1</sub>* | This study |
| p197_DG | *CEN, LEU2, P<sub>MSP1</sub>-MSP1-MSP1(EQ)-3XFLAG-T<sub>MSP1</sub>* | This study |
| p198_DG | *CEN, LEU2, P<sub>MSP1</sub>-MSP1(EQ)-MSP1(WT)-3XFLAG-T<sub>MSP1</sub>* | This study |
| p199_DG | *CEN, LEU2, P<sub>MSP1</sub>-MSP1(WT)-3XFLAG-T<sub>MSP1</sub>* | This study |
| p200_DG | *CEN, LEU2, P<sub>MSP1</sub>-MSP1(EQ)-3XFLAG-T<sub>MSP1</sub>* | This study |
| p202_DG | *CEN, HIS3, P<sub>TEF1</sub>-EGFP-T<sub>TEF1</sub>* | This study |
| p203_DG | *CEN, HIS3, P<sub>TEF1</sub>-mCherry-T<sub>TEF1</sub>* | This study |
| p204_DG | *CEN, HIS3, P<sub>TEF1</sub>-EGFP-E2A-mCherry-T<sub>TEF1</sub>* | This study |
| p205_DG | *CEN, HIS3, P<sub>TEF1</sub>-EGFP-Pex15ΔC30-E2A-mCherry-T<sub>TEF1</sub>* | This study |
| pHF_002 | His-TEV-Msp1 WT Monomer | Ref. <sup>35</sup> |
| pHF003 | His-TEV-Msp1 E193Q Monomer | Ref. <sup>35</sup> |
| pHF004 | His-TEV-Msp1 WT-WT Dimer | This study |
| pHF099 | His-TEV-Msp1 WT-EQ Dimer | This study |
| pHF129 | His-TEV-Msp1 EQ-WT Dimer | This study |
| pHF027 | GST-3C-SGTA | Ref. <sup>35</sup> |
| pHF050 | GST-3C-Calmodulin | Ref. <sup>35</sup> |
| p098_DG | His-MBP-Ubiquitin-mNG2 | Ref. <sup>35</sup> |
| p048_DG | His-3C-LgBiT | Ref. <sup>35</sup> |
| pHF122 | His-3C-Sumo-Sec22-HiBiT | Ref. <sup>35</sup> |

*cerevisiae* was judged by anti-Flag western blot with 1:10,000 dilution of rabbit anti-Flag antibody (Invitrogen) and a 1:10,000 dilution of goat anti-rabbit-HRP secondary antibody (Proteintech). Complementation was performed as described^17^. Briefly, haploid W303-1 cells containing the appropriate plasmid were grown overnight in SD -LEU with 2% glucose. Cultures were diluted to 0.02 OD_600_ in fresh SD -LEU media until mid-log phase. Cultures were washed 3x with sterile water and diluted to 1 OD_600_. Samples were serially diluted 5x and then spotted onto SD -LEU plates containing either 2% glucose or 2% glycerol and grown at 30° C. Images are representative of N > 2 trials.

### Flow cytometry assay

GFP-Pex15*Δ*C30-E2A-mCherry was cloned into a *CEN* and *HIS3* based vector and transformed into *Msp1Δ, Get3Δ* W303-1 yeast containing an Msp1 construct on a *CEN* and LEU2 based vector. Cells were grown in SD -HIS -LEU media at 30° C until it reached OD_600_ ∼ 3.0, then diluted to OD_600_ = 1 in sterile water media. Flow cytometry was completed on BD Biosciences LSR15 cytometer and further analysis completed using FlowJo as described previously^34^.

## Acknowledgements

The authors wish to thank members of the Wohlever lab for helpful discussions and feedback on the project.

## Funding

This work was supported by NIH grant R35GM137904 (MLW).

### Author Contributions

Conceptualization: BS, DG, MLW; Methodology: BS, DG, IW, NW, MLW; Investigation: BS, DG, IW, NW; Writing – Original Draft: BS, MLW; Writing – Review and Editing: BS, DG, IW, NW, MLW; Supervision and Funding Acquisition: MLW Competing Interests: The authors declare no competing interests.

## Notes

### Competing Interest Statement

The authors have declared no competing interest.

